# PopZ is dispensable for cell fitness but required for polar localization of ParB in *Zymomonas mobilis*

**DOI:** 10.64898/2026.07.24.740507

**Authors:** Katsuya Fuchino, Waldemar Vollmer, Richard Daniel

**Author notes:** Corresponding author Katsuya Fuchino.

## Abstract

Chromosome segregation is an essential process in monoploid bacteria, in which faithful inheritance of the genetic material is required for viability of daughter cell. In several alpha-proteobacteria, the polar organizing protein PopZ plays a crucial role in chromosome segregation by anchoring the chromosome partitioning machinery at a cell pole. While this process has been studied in monoploid species, it remains unclear whether this type of segregation mechanisms play a significant role in bacterial cells that carry multiple copies of chromosome.

The alpha-proteobacterium *Zymomonas mobilis* exhibits ethanologenic physiology, and thus, is a promising chassis for biofuel production. Recent studies showed that *Z. mobilis* may serve as a non-model system for studying bacterial cell biology, owing to its relatively simple physiology and reduced genome size. *Z. mobilis* has previously been shown to be polyploid, and possesses a homolog of PopZ, a key regulator of chromosome organisation and cell polarity in alpha-proteobacteria. This raises the question of whether PopZ is involved in the organization of multiple chromosome copies. Here, we investigated the function of PopZ in *Z. mobilis*. Unlike in previously studied alpha-proteobacterial species, PopZ is dispensable for growth and cell morphology in *Z. mobilis*. However, the loss of PopZ altered the polar accumulation of chromosome segregation protein ParB, indicating an involvement in chromosome organization in *Z. mobilis*. These findings indicate that *Z. mobilis* might possess a chromosome segregation machinery that is dispensable under favourable growth conditions but might become important under certain conditions that reduce chomorome copy numbers.

## Introduction

Inheritance of genetic information is essential for all life. Bacterial cells may be considered structurally simple compared to eukaryotic cells, but they possess sophisticated mechanisms to ensure replication of genetic material and its transmission to the daughter cells, which are well coordinated with cell growth and division. To date, our understanding of bacterial chromosome segregation has been studied mainly in monoploid model organisms, such as *Escherichia coli, Bacillus subtilis*, and *Caulobacter crescentus*. In these species, the machinery governing the physical separation of replicated chromosome is semi-essential because its absence resulting in anucleate cells or obstruction of cell division process.

Other bacterial species are known to possess two distinct chromosomes within a cell, such as *Rhodobacter sphaeroides* and *Vibrio cholerae*, and possess mechanisms to ensure both chromosome are located in each daughter cells (1, 2). Some bacteria exhibit multiple copies of chromosome within a cell (polyploidy). These species include cyanobacterial species, *Azotobacter vinelandii* and the gigantic bacteria *Epulopiscium* sp., and several other bacterial species. (3-6). Currently, knowledge about how these bacteria spatially organise its multiple copies of chromosome and partition chromosomes into daughter cells are limited.

Alpha-proteobacteria exhibits remarkable diversity in cell morphology, making them attracting living-system to study fundamental bacterial cell organisation (7). In this bacterial group, **<u>p</u>**olar **<u>o</u>**rganising **<u>p</u>**rotein PopZ has been shown to play fundamental processes such as chromosome segregation and the establishment of cell polarity (8). PopZ was discovered in the asymmetric cell model bacterium *C. crescentus*, and shown to play an important role in chromosome segregation by anchoring the chromosome partitioning ParB–*parS* complex to the cell pole, through interaction with the partitioning protein ParB which binds to the *parS* site in the chromosome. The loss of PopZ led to severe morphological defects that includes production of anucleated cells, mis-localised cell division, impaired growth and loss of the cell polarity. (9, 10) Owing to its physicochemical properties, PopZ appears to form condensate microdomains that act as an interaction hub at the cell pole, for protein partners which carry out important cellular processes and thereby establishes cell polarity (11-13). PopZ is widely conserved in alpha-proteobacteria, and its homologue was studied in *Agrobacterium tumefaciens* and *Magnetospirillum magneticum*, in which deletion of *popZ* caused severe cellular phenotypes (14-16).

Interestingly, PopZ homologue is conserved in a polyploid alpha-proteobacterium, *Zymomonas mobilis*. It is well known for its industrial use in biotechnology and alcohol beverage industry (17, 18). Previously, *Z. mobilis* has been shown to be polyploid (19-21). Thus, we were intrigued by the fundamental question, what biological role PopZ plays in *Z. mobilis* cells. Here, we investigated the function of PopZ in *Z. mobilis* and found that PopZ is dispensable for cell growth and division under laboratory conditions. Yet, PopZ was found to be required for polar accumulation of partitioning protein ParB and appears to contribute to chromosome organization.

## Methods

### Bacterial strains, plasmids, and growth conditions

Bacterial strains and plasmids used in this study are listed in Table S1. *Z. mobilis* was grown in RM complex medium containing glucose (20 g/L), yeast extract (5 g/L), NH_4_SO_4_ (1 g/L), KH_2_PO_4_ (1 g/L), and MgSO_4_ (0.5 g/L). As *Z. mobilis* is a facultative anaerobe, the complex medium was briefly sparged with nitrogen gas to make it micro-aerobic. For genetic manipulation, 12 mL cultures of *Z. mobilis* were incubated at 30°C in a capped 15 mL Falcon tube without shaking. For obtaining the growth profiles, the optical density was measured in a 96-well plate using SPECTROstar Nano (BMG labtech).

### Genetic manipulation of *Z. mobilis* strains

We employed a published method for genetic manipulations (21). To construct plasmids for genetic manipulation, the upstream and downstream of the gene of interest (500 bp) were amplified by polymerase chain reaction (PCR) using Q5 DNA polymerase (New England Biolabs). The obtained fragments were inserted into the plasmid pPK15534 by Gibson Assembly (New England Biolabs). The sequence of constructed plasmids was verified by Sanger sequencing or whole plasmid sequencing (Eurofins Genomics). The plasmids were transferred into *Z. mobilis* strain ZM6 via conjugation with *Escherichia coli* strain WM6026 as described (22). *Z. mobilis* cells with the integrated plasmid in their chromosomal DNA were selected by 90 μg/mL chloramphenicol in RM medium agar. The integration of the plasmid into the chromosomal DNA was confirmed by colony PCR. For excision of the plasmid, the *Z. mobilis* cells were grown overnight in liquid RM medium without selection, and subsequently the loss of plasmid was achieved by counter-selection with the *Bacillus subtilis* SacB that had been optimised for *Z. mobilis* (21).

For cloning of P_*pdc*_-*popZ* into the replicative plasmid pBBR, the *popZ* gene (*ZZ6_1271*) and the promoter region of *pdc* were amplified by Q5 DNA polymerase. These two fragments were annealed by overlap PCR using Q5 DNA polymerase, and the annealed product and the plasmid pBBR were cut by the restriction enzyme XbaI and XhoI, and washed using a PCR purification kit (Qiagen). These two products were ligated by T4 ligase (New England Biolabs) and transferred into *Z. mobilis* strains via conjugation. Ex-conjugants were selected on RM medium containing kanamycin (220 μg/mL). The primers used for constructing and verifying the strains are listed in Table S2.

### Microscopy and image analysis

A culture with growing *Z. mobilis* (0.5 µl) was spotted onto a 1% agarose-pad made of RM growth medium and covered by a cover-glass. Phase contrast and fluorescence images of *Z. mobilis* cells were taken using a Nikon Eclipse Ti microscope (Nikon) equipped with a Plan Apochromat 100x objective with an NA of 1.40 and Ph3 Phase plate (Nikon) and a Prime sCMOS camera (Teledyne Photometrics). HADA and ParB-sfGFP was imaged using 49000 and 49002 filter cubes respectively (Chroma). Images were acquired using Nikon NIS elements AR software. Image analysis was performed using MicrobeJ (23). Unpaired T-test was applied to evaluate statistical significance. For visualizing the PG growth sites in *Z. mobilis*, cells were grown for 6 hours in the growth medium containing the fluorescent D-amino acid HADA (7-hydroxycoumarincarbonylamino-D-alanine) (24) at a final concentration of 500 μM. After the growth, HADA was removed by washing the cells in PBS before they were spotted on the RM agarose-pad. Phase contrast and fluorescence images were taken immediately after the spotting, and the cells were incubated on the RM medium at room temperature for 55 minutes before images were taken again. Demographs of the fluorescence signal were obtained using MicrobeJ (23).

### Bacterial two-hybrid assay

For cloning of *parB* and *popZ* into the plasmid pUT18/18C and pKNT25, the *popZ* gene (*ZZ6_1271*) was amplified by Q5 DNA polymerase. The PCR products and plasmids pUT18C and pKNT25 were cut by the restriction enzyme XbaI and KpnI. pUT18 and its insert were cut by XbaI and HindIII. The cut products were washed using a PCR purification kit (Qiagen), and ligated by T4 ligase (New England Biolabs). The sequence of constructed plasmids was verified by Sanger sequencing or whole plasmid sequencing (Eurofins Genomics). The constructed plasmids were introduced into *E. coli* strain BTH101 by chemical transformation. Bacterial two-hybrid assay was performed as described in (25). As negative controls, *E. coli strain* BTH carrying pKNT25-empty and pUT18/18C-empty were used in the assay. As a positive control, the strain carrying pKNT25-*zip* and pUT18C-*zip* was included. The *E. coli* strains were grown on LB agar plate supplemented with X-gal (40 μg/ml) and IPTG (0.5 mM) for 24 hours at 30 °C, and the images were taken.

## Results

### Lack of PopZ does not impair growth and morphology in *Z. mobilis*

We previously visualized approximate cellular locations of the *oriC* site of the chromosomes using ParB-sfGFP labelling. We found that several ParB-sfGFP foci were distributed throughout the cell, with an accumulated focus located at one of the cell poles, mostly at the old cell pole (21). This observation raised the question if the multi-copied chromosomes are spatially controlled within a *Z. mobilis* cell and, if so, whether this control is important for cell fitness. To address this question, we investigated the role of the polar organizing protein PopZ homologue in *Z. mobilis*. We first generated a *popZ* deletion mutant (Δ*popZ*), using a previously developed homologous-recombination based method (21). We confirmed that *popZ* was deleted in all copies of the chromosome and then compared the growth of the mutant and that of its parental strain ZM6 in the RM growth medium supplemented with 2% glucose. This analysis showed that Δ*popZ* appears to exhibit no obvious growth or morphological phenotype under regular growth conditions (Fig 1AB). These observations are in sharp contrast to the growth defects reported in other alpha-proteobacteria (9, 14, 16). We also examined growth of the mutant in the presence of elevated salt concentrations. *Z. mobilis* cells are sensitive to salt, even at the mildly elevated concentration of 0.225 M NaCl, producing elongated cells with one bulged cell pole and limited formation of septa (26). Interestingly, Δ*popZ* exhibited no observable growth or morphological defects under salt conditions (Fig. S1), showing that PopZ is dispensable for cell growth in *Z. mobilis*.

**Fig. 1.**
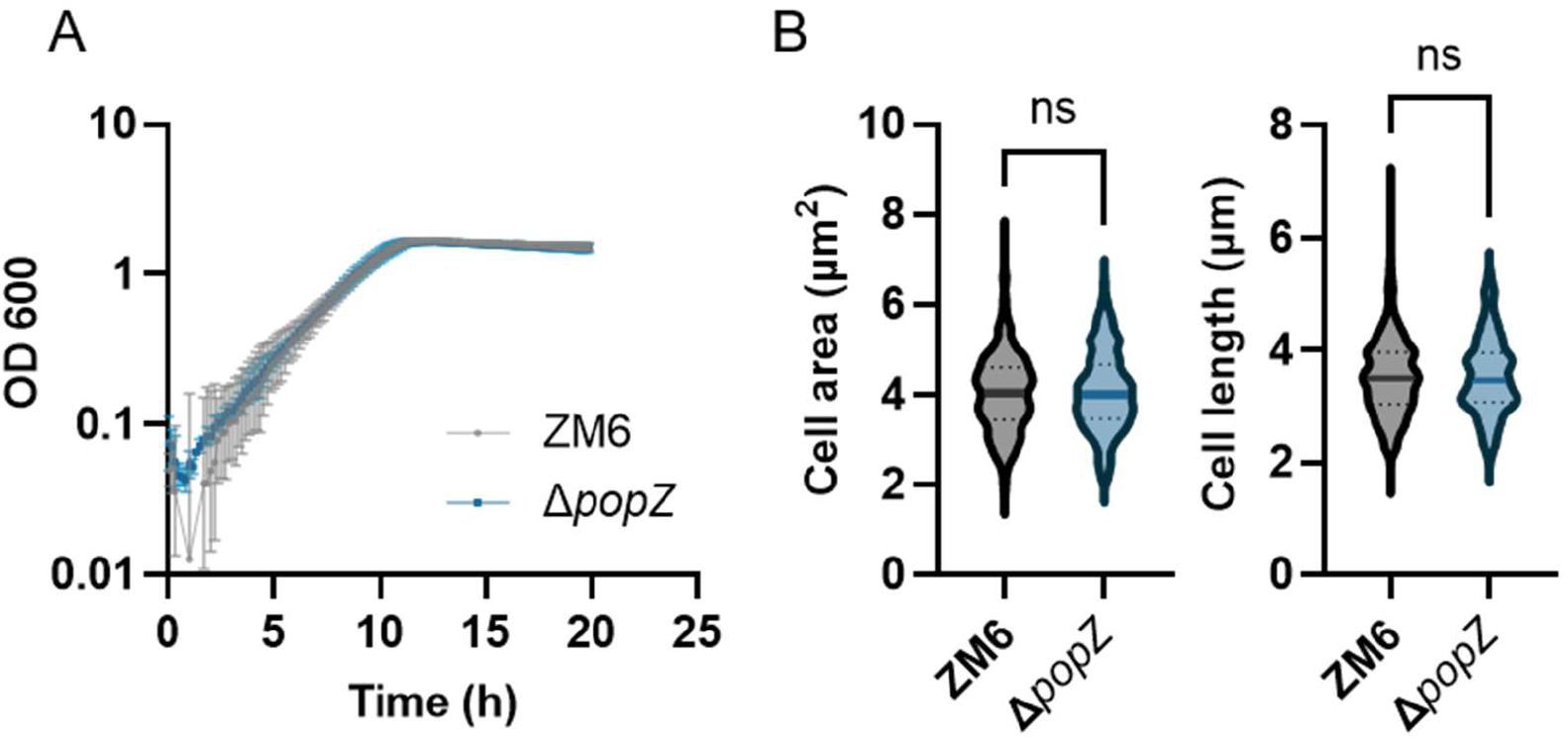
Loss of *popZ* did not influence cell growth and morphology in *Z. mobilis*. **(A)** Growth of *Z. mobilis* strains ZM6 (wild-type) and Δ*popZ* under regular growth conditions (RM medium with 2% glucose), measured by a 96-well plate reader. Biological replicates N = 3. The error bars in the plot represent the standard deviation. **(B)** Box-and-whisker (violin) plot presenting the cell area and length of *Z. mobilis* cells of the two strains grown to an OD600 of 0.6-1.3. Sample size, n = 434 cells (ZM6), n = 317 cells (Δ*popZ*). ns, not significant.

### ParB-sfGFP loses polar localisation in the *popZ* mutant

PopZ was previously shown to interact with ParB, and deletion of *popZ* caused mis-localisation of ParB in *C. crescentus* (10) and *A. tumefaciens* (14). We therefore aimed to assess whether the absence of PopZ also impairs ParB localization in *Z. mobilis*. For this, we deleted *popZ* in a strain expressing *parB-sfGFP* at its native locus, generating Δ*popZ* + *parB*-*sfGFP*. We examined the subcellular localization of sfGFP signals and compared them to that of parental strain.

From the fluorescence images it was seen that the amount of ParB-sfGFP localising to the cell pole in the Δ*popZ* was significantly reduced relative to that observed in ZM6 (Fig. 2A and S2). Quantification of the signals showed that the mean value of sfGFP signal from an accumulated, polar focus was 16053 in the wild-type, and it was reduced to 8781 in Δ*popZ* (Fig. 2B). Furthermore, while more than half of the ZM6 + *parB*-*sfGFP* cells in the population (53.5%, N = 142) exhibited an accumulated polar signal which was at least 50% stronger than that of the second-intensified focus within the same cell, the Δ*popZ* + *parB*-*sfGFP* cells only exhibited 8.6 % of such foci (N = 160). To confirm that this was not due to polar effects of the genetic manipulation, we inserted *popZ* with its natural promoter at a different locus (between ZZ6_1059 and ZZ6_1060) in the chromosome of Δ*popZ* + *parB*-*sfGFP* (trans-complementation). The complementation restored the intensity of the ParB-sfGFP signal at the pole (Fig. 2B) and its pronounced accumulation was found in 56% of cells in the population (N = 182). These data confirmed that the loss of PopZ led to the reduced accumulation of ParB at the cell pole.

**Fig. 2.**
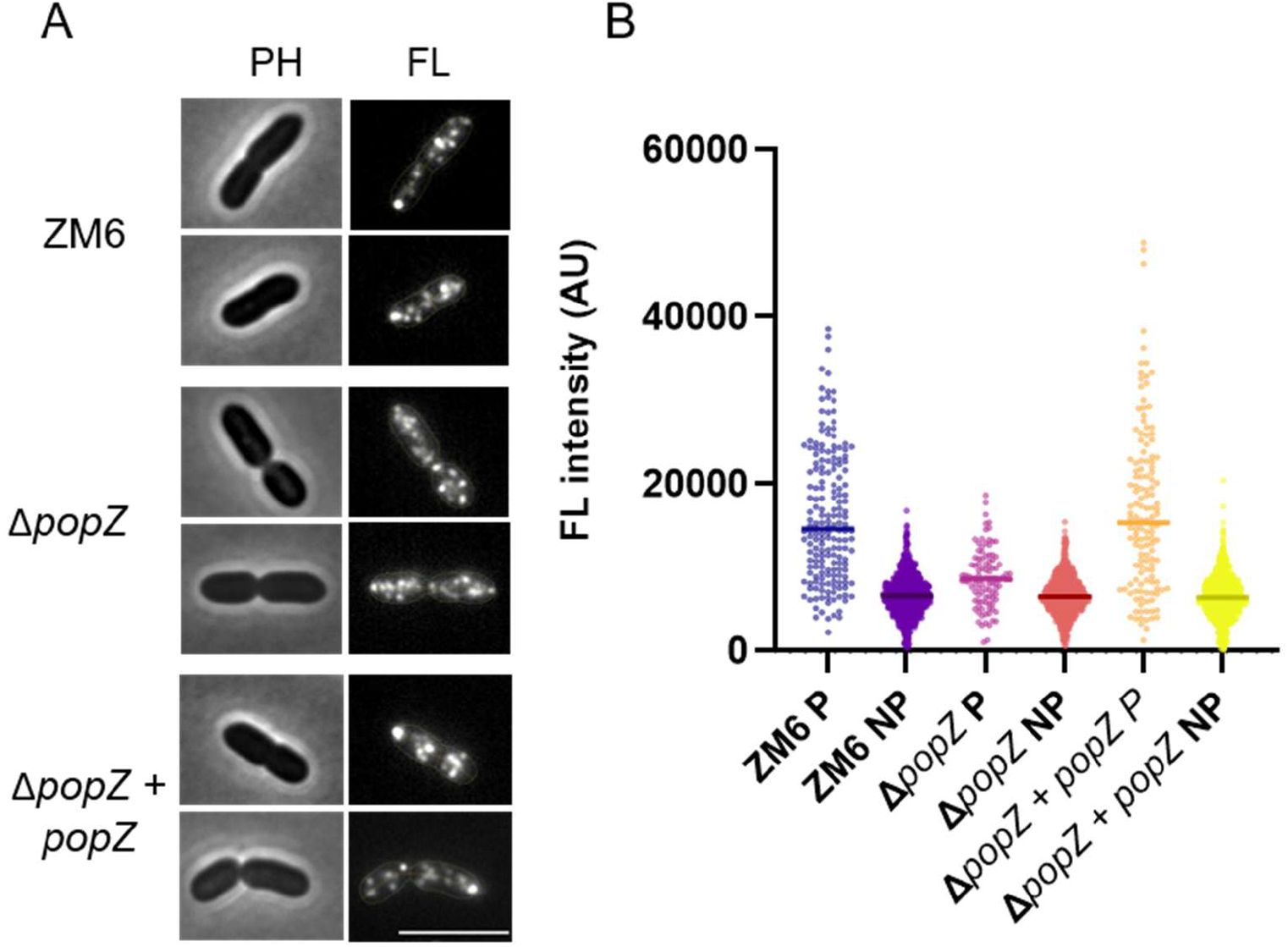
ParB polar-localisation is altered in the *popZ* mutant. **(A)** Phase contrast (left) and fluorescence (right) images of growing ZM6 (wild-type) + *parB*-*sfGFP*, Δ*popZ* + *parB*-*sfGFP*, Δ*popZ* + *popZ* (complementation strain) + *parB*-*sfGFP* cells under regular growth conditions. Contour of cells in the fluorescence images were added by MicrobeJ (23). Scale bar, 5 μm. For simplicity, ‘+ *parB*-*sfGFP*’ was omitted from the strain names in the figure. **(B)** A plot presenting ParB-sfGFP fluorescence intensity of polar focus (indicated by P) and foci observed from other cell regions (indicated by NP). *Z. mobilis* cells of the three different strains were grown to an OD600 of 0.7-1.5. Sample size, n = 180 foci (ZM6 P), n = 929 foci (ZM6 NP) n = 104 (Δ*popZ* P) n = 959 (Δ*popZ* NP) n = 136 foci (Δ*popZ* + *popZ* P) n = 882 (Δ*popZ* + *popZ* NP). Unpaired T-test was applied between strains, showing the significant difference between ZM6 P and Δ*popZ* P (P < 0.0001). No significant difference was found between ZM6 P and Δ*popZ* + *popZ* P (P = 0.4269), ZM6 NP and Δ*popZ* NP (P = 0.5792), ZM6 NP and Δ*popZ* + *popZ* NP (P = 0.0932). For simplicity, ‘+ *parB*-*sfGFP*’ was omitted from the strain names in the figure.

To further characterise this effect, we performed time-lapse imaging on both wild-type and Δ*popZ* carrying the ParB-sfGFP fusion. In the wild-type background, accumulated signals at one of the cell poles stayed in most of the cells during growth, and the several foci moved around in the cytoplasm (Fig. S3). In the Δ*popZ* background, the ParB-sfGFP foci rarely accumulated at a cell pole, but the movement of the foci occurred with similar dynamics as in cells with PopZ (Fig. S3). Thus, PopZ likely anchors ParB at the pole, as previously shown in other studied bacteria.

To further understand biological function of PopZ in *Z. mobilis* cells, we overproduced PopZ and assessed the effect on cell fitness and ParB localisation. We constructed *popZ* under the constitutively active *pdc* promoter (P_*pdc*_) in the replicative plasmid pBBR, and introduced it into *Z. mobilis*. This promoter has been shown to be highly active in *Z. mobilis* (27, 28). Interestingly, the overexpression of PopZ did not cause any particular growth defects or change in cell morphology (Fig. 3AB), in sharp contrast to other alpha-proteobacteria where cellular organisation was severely influenced by the overexpression of the *popZ* homologue (9, 14). Also, introduction of this overexpression construct into the ParB-sfGFP strain gave no indication of a change in the overall localization pattern of ParB. (Fig. 3CD). Furthermore, in this overexpression background phase-bright focus was occasionally observed near at a cell pole within the cells (Fig. S4). These were not evident in the wild-type background carrying empty plasmid. These foci might reflect the phase-separation / condensate property of accumulated PopZ in the cytoplasm. Overexpression of *popZ* in other studied species resulted in the accumulation of PopZ near the cell pole, which excluded macromolecules from the accumulated region (9, 16). Similar phenomenon might have occurred in *Z. mobilis* cells overproducing PopZ.

**Fig. 3.**
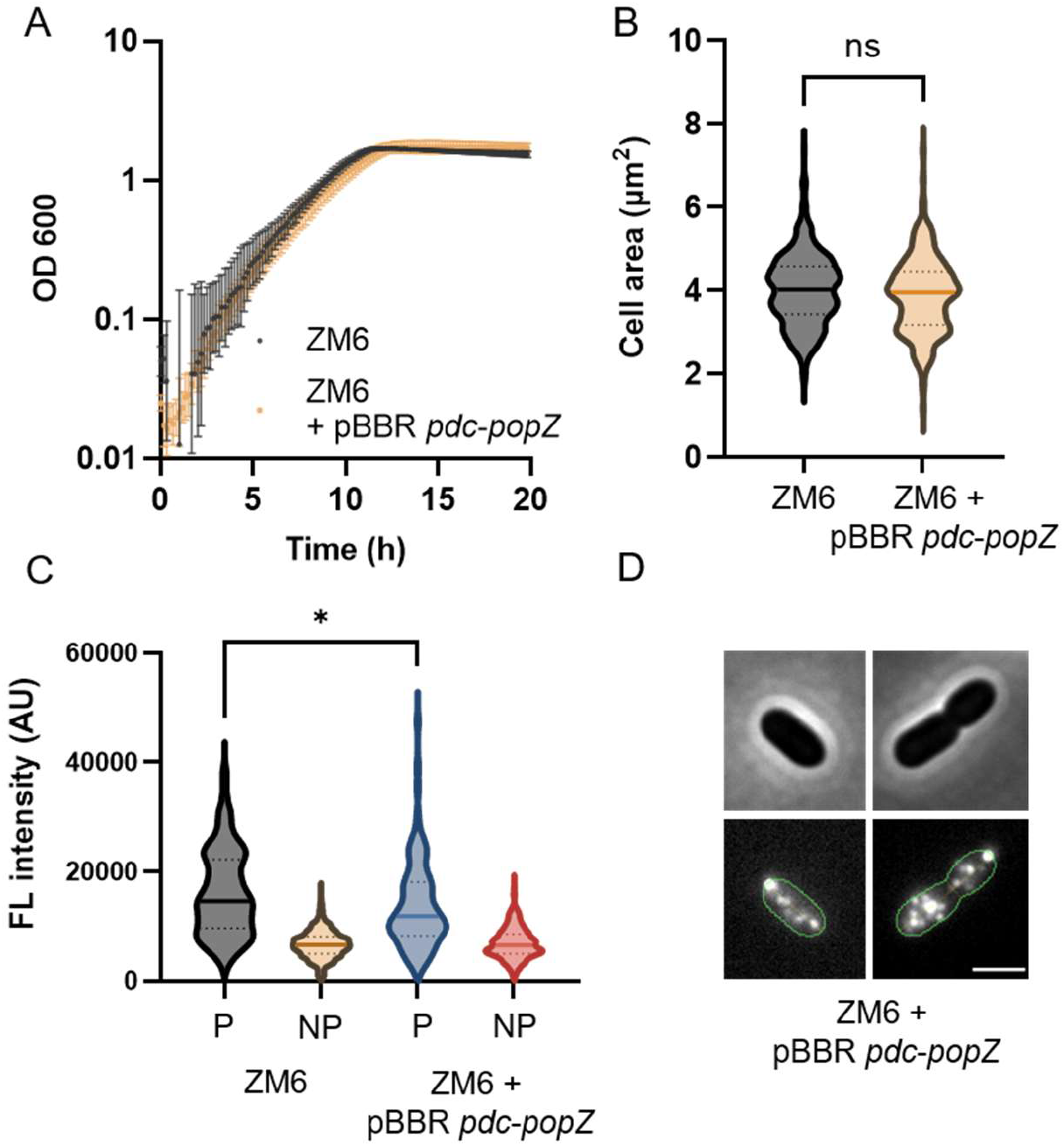
Overexpression of *popZ* does not impair growth and ParB localisation. **(A)** Growth of *Z. mobilis* strains ZM6 (wild-type) + *parB*-*sfGFP*, and ZM6 + *parB*-*sfGFP* + pBBR P_*pdc*_-*popZ* under regular growth conditions. **(B)** Box-and-whisker (violin) plot presenting the cell area of growing *Z. mobilis* cells of the two strains. Sample size, n = 434 cells (ZM6 + *parB*-*sfGFP*), n = 241 cells (ZM6 + *parB*-*sfGFP* + pBBR P_*pdc*_-*popZ*), ns, not significant. **(C)** Box-and-whisker (violin) plot presenting ParB-sfGFP fluorescence intensity of polar focus (indicated by P) and foci observed from other cell regions (indicated by NP). Sample size, n = 180 foci (ZM6 + *parB*-*sfGFP* P), n = 928 foci (ZM6 + *parB*-*sfGFP* NP) n = 97 (ZM6 + *parB*-*sfGFP* + pBBR P_*pdc*_-*popZ* P) n = 694 foci (ZM6 + *parB*-*sfGFP* pBBR P_*pdc*_-*popZ* NP). *, P = 0.0255 (unpaired t-test). **(D)** Phase contrast (top) and fluorescence (bottom) images of growing ZM6 + *parB*-*sfGFP* carrying pBBR P_*pdc*_-*popZ* under regular growth conditions (OD600 of 0.5 -0.9). Contour of cells in the fluorescence images were added by MicrobeJ (23). Scale bar, 2 μm. For simplicity, ‘+ *parB*-*sfGFP*’ was omitted from the strain names in the figure.

### PopZ interacts with ParB in *Z. mobilis*

Based on the obtained results, we speculated that PopZ localises at the cell pole, as in other studied bacteria, and interacts with ParB. For imaging the localisation, we fused PopZ with a fluorescent protein mCherry at its C-terminus. However, the fusion protein did not complement the Δ*popZ* mutant phenotype (ParB-sfGFP localisation) when introduced at the locus used for the complementation, indicating that the fusion was not functional.

To assess the interaction between ParB and PopZ, We employed a bacterial two-hybrid assay that detects protein interactions through reconstituting adenylate cyclase activity in the *Escherichia coli* cytoplasm (25). The *parB* gene was cloned into the pUT18 and pUT18C plasmid that carries the T18 fragment of adenylate cyclase, and the *popZ* was cloned into the pKNT25 plasmid carrying the T25 fragment. Although a false-positive was observed with one of the negative control constructs, ParB and PopZ appeared to interact in the assay, as indicated by the blue colouration of the colony reconstituting adenylate cyclase activity, presumably through the interaction between ParB and PopZ in the *E. coli* cytoplasm (Fig. S5). Nevertheless, this is a preliminary result and additional analyses will be required to confirm the interaction.

### *Z. mobilis* cells grow predominantly around mid-cell

*Z. mobilis* ParB polar accumulation predominantly occurred at an old cell pole of dividing cells (Fig. S2 and S3). It is likely that PopZ recruits ParB to the pole, but the basis for PopZ polar localisation remained unclear. In addition, *Z. mobilis* cells do not exhibit obvious cell polarity like the asymmetric *C. crescentus* cells or polarly growing *A. tumefaciens* cells. A recent cell biological study, using fluorescent D-amino acids (24) that can be incorporated into the PG, revealed that growth patterns even among *Caulobacteraceae* family of alphaproteobacterial are diverse. Some species like *Asticcacaulis excentricus* grow unidirectionally at the mid-cell towards new pole while *C. crescentus* exhibits bidirectional growth (29). A previous study using fluorescent D-amino acids (HADA) showed that newly synthesised materials were visualised around the mid-cell and lateral wall regions in *Z. mobilis*, (30). However, it remained unclear whether growth occurs in a specific direction. If particular directional growth does occur in *Z. mobilis*, it may be related to the biased polar localization of ParB and, presumably, PopZ, in the daughter cell after cell separation. Thus, we pulse-chased growing *Z. mobilis* cells with HADA and monitored the disappearance of the fluorescence signal during growth, following an approach similar to that performed in (29). After the cells were grown and stained with HADA for about 6 hours (about 4 generation) under regular growth conditions, the cells were washed and spotted on the RM agar-pad to monitor the signals. Immediately after the imaging started (time = 0), the signal was detected throughout the cell’s periphery, with pronounced accumulation at the septum (Fig. 4A). After growth for 55 mins, most of the cells exhibited a reduced HADA signal near mid-cell (Fig. 4B). Furthermore, the loss of fluorescence signal extended bidirectionally from the septum (Fig. 4B). These unlabelled regions indicate that the cells incorporated new PG around the mid-cell regions toward both poles. In a complementary approach, we stained the growing cells with HADA for 20 mins and observed the signal at new grown sites. The signal was enhanced around the mid-cell region and the septum (Fig. S6), consistent with the results from the pulse-chase experiment. Importantly, the staining was also observed in cells that did not form a septum yet (Fig. S6). These data suggest that *Z. mobilis* cells grow predominantly around the mid-cell regions, and do not exhibit a unidirectional growth. Thus, the asymmetric localization of ParB at one cell pole is not mediated by biased cell growth.

**Fig. 4.**
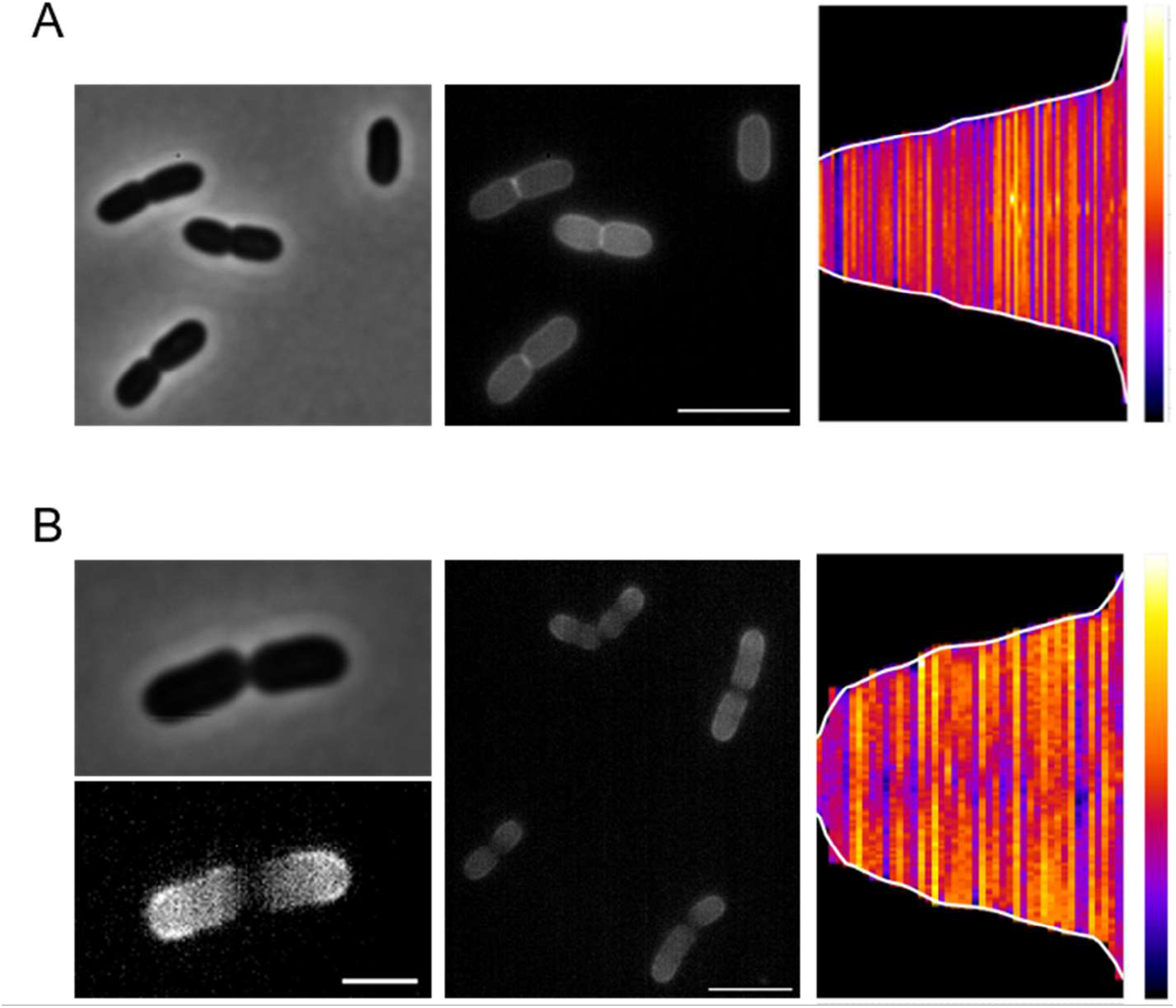
*Z. mobilis* predominantly grows around mid-cell position through a bidirectional manner. **(A)** Representative phase contrast (left) and fluorescence (middle) image of growing *Z. mobilis* cells stained by HADA for 4 generation time. HADA was removed by washing before imaging before being spotted on the RM agarose-pad. Scale bar, 5 μm. (Right): A demograph showing the localization and intensities of HADA fluorescence signal from individual cells in the population. The fluorescence intensity ranged from 4,000 to 9,000, corresponding to the color scale from dark blue to yellow. N = 75. Biological replicates = 2 **(B)** The spotted cells were grown on the agar pad for 55 mins before imaging. (Left): Representative images (top: phase contrast, bottom: fluorescence) of the stained *Z. mobilis* cell grown on the agar pad for 55 mins before imaging. Scale bar, 2 μm. (Middle): a fluorescence image of other stained *Z. mobilis* cells grown on the agar pad for 55 mins. Scale bar, 5 μm. (Right): A demograph showing the localization and intensities of HADA fluorescence signal from individual cells in the population. The fluorescence intensity ranged from 3,800 to 5,700, corresponding to the color scale from dark blue to yellow. N = 75. Biological replicates = 2

## Discussion

PopZ was dispensable and its overexpression did not cause detectable cellular malfunction in *Z. mobilis* under regular growth conditions, in contrast to previously studied bacteria. Although our data suggest that PopZ influences the localization of the chromosome partitioning protein ParB, possibly through a protein-interaction (Fig. 2 and Fig. S5), this was not required for cell fitness under the conditions examined. As *Z. mobilis* ZM6 cells have previously been shown to contain approximately 5–20 chromosome copies (20, 21), chromosomes may be delivered into daughter cells through a passive process, such as diffusion of chromosomes within the cytoplasm, even in the absence of an active segregation mechanism. As alpha-proteobacteria exhibit remarkable diversity in cell organization, it would be of considerable interest to investigate the function of PopZ in a broader range of species of this phylum to understand how PopZ has adapted to the biology of each species.

Interestingly, *Z. mobilis* cells divide around the mid-cell region (30). This mode of division might also reflect its polyploidy. Precise mid-cell division is important in monoploid bacteria to ensure chromosome inheritance. In contrast, accurate mid-cell division may be less important in polyploid cells, because multiple copies of the chromosome may reduce the risk of producing anucleate daughter cells. Yet, cell division is still biased toward mid-cell, indicating that a mechanism exists to position the division machinery. It would be interesting to reveal the basis of this positioning mechanism, and whether it is linked to chromosome.

ParB-sfGFP mostly remained monopolar throughout the cell cycle in *Z. mobilis* (Fig. S3). This raises a possibility that PopZ is likewise restricted to a single cell pole. PopZ may function to retain the chromosome at one specific cell pole rather than to capture the newly segregated ParB-complex at the opposite pole, as observed in other species. The accumulation of ParB-sfGFP signal at the cell pole indicates that chromosome replication likely occurs at this site. Thus, utilizing its intrinsic property to localize at the pole, PopZ may contribute to spatial organization of replication. However, testing this hypothesis will require determining the localisation of PopZ and the proteins involved in the replication machinery simultaneously. In addition, it is currently not clear at which cell pole ParB localizes in the daughter cell that did not inherit the accumulated ParB-sfGFP focus. We were unable to clarify this from our time-lapse imaging because a prolonged imaging caused phototoxicity. Resolving localization pattern of polar accumulation of ParB over several generations should provide further insight into the function of ParB and PopZ.

What is the advantage of possessing PopZ in *Z. mobilis*? A previous study showed that *Z. mobilis* cells were able to grow for several generations in the absence of environmental phosphorous, during which its chromosome copy number per single cell significantly decreased (31). Cell growth and division without essential phosphorous nutrient is likely mediated by passing the multiple copies of chromosomes into daughter cells. DNA might be used as a phosphorous source through its degradation as well. It is tempting to speculate that PopZ and other machinery involved in the spatial control of the chromosomes become critical for cell fitness when the copy number decreases. We attempted to test this hypothesis, but the strain ZM6 was not able to grow in the synthetic medium used for the strain ZM4 (31, 32), making this experiment technically infeasible. We are currently searching for a suitable synthetic growth medium to enable the experiment.

## Acknowledgements

This work was supported by Newton International Fellowship (The Royal Society) to K.F. (NIF\R1\221190) and BBSRC Research Grant to R.D. (BB/Y002644/1).

## References

1. Dubarry N, Willis CR, Ball G, Lesterlin C, Armitage JP. 2019. In Vivo Imaging of the Segregation of the 2 Chromosomes and the Cell Division Proteins of Rhodobacter sphaeroides Reveals an Unexpected Role for MipZ. mBio 10:10.1128/mbio.02515-18.

2. Espinosa E, Challita J, Desfontaines J-M, Possoz C, Val M-E, Mazel D, Marbouty M, Koszul R, Galli E, Barre F-X. 2024. MatP local enrichment delays segregation independently of tetramer formation and septal anchoring in Vibrio cholerae. Nature Communications 15:9893.

3. Mendell JE, Clements KD, Choat JH, Angert ER. 2008. Extreme polyploidy in a large bacterium. Proceedings of the National Academy of Sciences 105:6730–6734.

4. Soppa J. 2014. Polyploidy in archaea and bacteria: about desiccation resistance, giant cell size, long-term survival, enforcement by a eukaryotic host and additional aspects. J Mol Microbiol Biotechnol 24:409–19.

5. Jain IH, Vijayan V, O’Shea EK. 2012. Spatial ordering of chromosomes enhances the fidelity of chromosome partitioning in cyanobacteria. Proceedings of the National Academy of Sciences 109:13638–13643.

6. Maldonado R, Jiménez J, Casadesús J. 1994. Changes of ploidy during the Azotobacter vinelandii growth cycle. Journal of Bacteriology 176:3911–3919.

7. Randich AM, Brun YV. 2015. Molecular mechanisms for the evolution of bacterial morphologies and growth modes. Frontiers in Microbiology Volume 6–2015.

8. Bergé M, Viollier PH. 2018. End-in-Sight: Cell Polarization by the Polygamic Organizer PopZ. Trends in Microbiology 26:363–375.

9. Ebersbach G, Briegel A, Jensen GJ, Jacobs-Wagner C. 2008. A Self-Associating Protein Critical for Chromosome Attachment, Division, and Polar Organization in Caulobacter. Cell 134:956–968.

10. Bowman GR, Comolli LR, Zhu J, Eckart M, Koenig M, Downing KH, Moerner WE, Earnest T, Shapiro L. 2008. A Polymeric Protein Anchors the Chromosomal Origin/ParB Complex at a Bacterial Cell Pole. Cell 134:945–955.

11. Holmes JA, Follett SE, Wang H, Meadows CP, Varga K, Bowman GR. 2016. Caulobacter PopZ forms an intrinsically disordered hub in organizing bacterial cell poles. Proceedings of the National Academy of Sciences 113:12490–12495.

12. Lasker K, Boeynaems S, Lam V, Scholl D, Stainton E, Briner A, Jacquemyn M, Daelemans D, Deniz A, Villa E, Holehouse AS, Gitler AD, Shapiro L. 2022. The material properties of a bacterial-derived biomolecular condensate tune biological function in natural and synthetic systems. Nat Commun 13:5643.

13. Nordyke CT, Ahmed YM, Puterbaugh RZ, Bowman GR, Varga K. 2020. Intrinsically Disordered Bacterial Polar Organizing Protein Z, PopZ, Interacts with Protein Binding Partners Through an N-terminal Molecular Recognition Feature. Journal of Molecular Biology 432:6092–6107.

14. Ehrle Haley M, Guidry Jacob T, Iacovetto R, Salisbury Anne K, Sandidge DJ, Bowman Grant R. 2017. Polar Organizing Protein PopZ Is Required for Chromosome Segregation in Agrobacterium tumefaciens. Journal of Bacteriology 199:10.1128/jb.00111-17.

15. Howell M, Aliashkevich A, Salisbury AK, Cava F, Bowman GR, Brown PJB. 2017. Absence of the Polar Organizing Protein PopZ Results in Reduced and Asymmetric Cell Division in Agrobacterium tumefaciens. Journal of Bacteriology 199:10.1128/jb.00101-17.

16. Pfeiffer D, Toro-Nahuelpan M, Bramkamp M, Plitzko JM, Schüler D. 2019. The Polar Organizing Protein PopZ Is Fundamental for Proper Cell Division and Segregation of Cellular Content in Magnetospirillum gryphiswaldense. mBio 10:10.1128/mbio.02716-18.

17. Braga A, Gomes D, Rainha J, Amorim C, Cardoso BB, Gudiña EJ, Silvério SC, Rodrigues JL, Rodrigues LR. 2021. Zymomonas mobilis as an emerging biotechnological chassis for the production of industrially relevant compounds. Bioresources and Bioprocessing 8:128.

18. Yang S, Fei Q, Zhang Y, Contreras LM, Utturkar SM, Brown SD, Himmel ME, Zhang M. 2016. Zymomonas mobilis as a model system for production of biofuels and biochemicals. Microbial Biotechnology 9:699–717.

19. Brenac L, Baidoo EEK, Keasling JD, Budin I. 2019. Distinct functional roles for hopanoid composition in the chemical tolerance of Zymomonas mobilis. Molecular Microbiology 112:1564–1575.

20. Fuchino K, Wasser D, Soppa J. 2021. Genome Copy Number Quantification Revealed That the Ethanologenic Alpha-Proteobacterium Zymomonas mobilis Is Polyploid. Frontiers in Microbiology Volume 12 - 2021.

21. Fuchino K, Astraios C, Daniel R, Grimshaw J, Banzhaf M, Vollmer W. 2026. Periplasmic SacB as a robust counter-selection tool for genome engineering in the polyploid bacterium Zymomonas mobilis. bioRxiv doi:10.64898/2026.01.13.699301:2026.01.13.699301.

22. Lal PB, Wells FM, Lyu Y, Ghosh IN, Landick R, Kiley PJ. 2019. A Markerless Method for Genome Engineering in Zymomonas mobilis ZM4. Frontiers in Microbiology Volume 10 - 2019.

23. Ducret A, Quardokus EM, Brun YV. 2016. MicrobeJ, a tool for high throughput bacterial cell detection and quantitative analysis. Nature Microbiology 1:16077.

24. Kuru E, Hughes HV, Brown PJ, Hall E, Tekkam S, Cava F, de Pedro MA, Brun YV, VanNieuwenhze MS. 2012. In Situ Probing of Newly Synthesized Peptidoglycan in Live Bacteria with Fluorescent D-Amino Acids. Angewandte Chemie International Edition 51:12519–12523.

25. Karimova G, Pidoux J, Ullmann A, Ladant D. 1998. A bacterial two-hybrid system based on a reconstituted signal transduction pathway. Proceedings of the National Academy of Sciences 95:5752–5756.

26. Fuchino K, Bruheim P. 2020. Increased salt tolerance in Zymomonas mobilis strain generated by adaptative evolution. Microbial Cell Factories 19:147.

27. Anggarini S, Murata M, Kido K, Kosaka T, Sootsuwan K, Thanonkeo P, Yamada M. 2020. Improvement of Thermotolerance of Zymomonas mobilis by Genes for Reactive Oxygen Species-Scavenging Enzymes and Heat Shock Proteins. Frontiers in Microbiology Volume 10 - 2019.

28. Fuchino K, Nalbant A, Gray J, Vollmer W. 2026. Peptidoglycan remodeling improves salt resilience of Zymomonas mobilis. Applied and Environmental Microbiology 92:e02350–25.

29. Delaby M, Yang L, Jacq M, Gallagher KA, Kysela DT, Hughes V, Pulido F, Veyrier FJ, VanNieuwenhze MS, Brun YV. 2025. Phenotypic plasticity in cell elongation among closely related bacterial species. Nature Communications 16:5099.

30. Fuchino K, Chan H, Hwang LC, Bruheim P. 2021. The Ethanologenic Bacterium Zymomonas mobilis Divides Asymmetrically and Exhibits Heterogeneity in DNA Content. Appl Environ Microbiol 87.

31. Brück P, Wasser D, Soppa J. 2023. One Advantage of Being Polyploid: Prokaryotes of Various Phylogenetic Groups Can Grow in the Absence of an Environmental Phosphate Source at the Expense of Their High Genome Copy Numbers. Microorganisms 11:2267.

32. Fuchino K, Kalnenieks U, Rutkis R, Grube M, Bruheim P. 2020. Metabolic Profiling of Glucose-Fed Metabolically Active Resting Zymomonas mobilis Strains. Metabolites 10:81.

