## Supplemental Figure S1-S6 and Supplemental table S1 -2 for "PopZ is dispensable for cell fitness but required for polar localization of ParB in *Zymomonas mobilis*"

**Supplemental Figures**

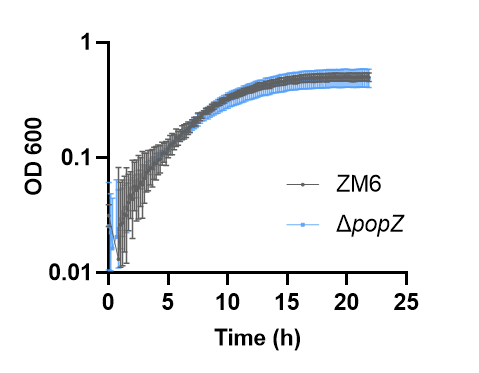

**Fig. S1** **Δ*popZ* did not exhibit increased sensitivity to salt stress compared with the wild-type.** Growth of *Z. mobilis* strains ZM6 (wild-type) and Δ*popZ* under salt stress conditions (NaCl 0.225 M). Biological replicates N = 3. The error bars represent the standard deviation.

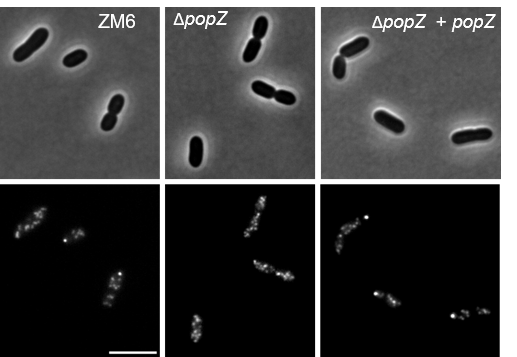

**Fig. S2 ParB-sfGFP localisation in *Z. mobilis* strains.**

(A) Phase contrast (top) and fluorescence (bottom) images of growing ZM6 (wild-type) + *parB*-*sfGFP*, Δ*popZ* + *parB*-*sfGFP*, Δ*popZ* + *popZ* (complementation strain) + *parB*-*sfGFP* cells under regular growth conditions. Scale bar, 5 μm.

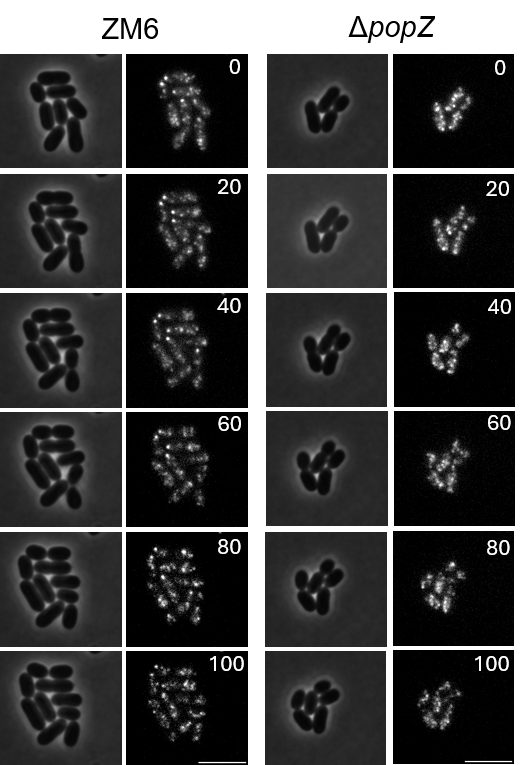

**Fig. S3 Time lapse imaging of ParB-sfGFP localisation in *ZM6* and Δ*popZ*.**

Time-lapse imaging of growing ZM6 + *parB-sfGFP* (left two panels) and Δ*popZ* + *parB-sfGFP* (right two panels) cells on 1% agarose-pad in the growth medium RM. Phase contrast (left) and fluorescence (right) images were taken every 10 min by fluorescence microscopy. Numbers in the images indicate time (minutes) after the imaging started. Scale bars, 10 μm.

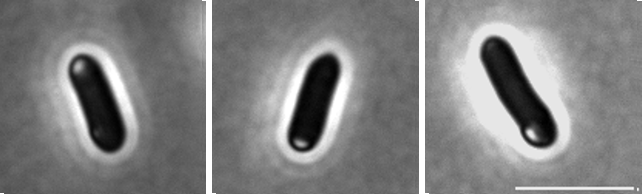

**Fig. S4** **Overexpression of *popZ* results in occasional formation of phase-bright structures.**

Representing phase contrast images of growing ZM6 + *parB*-*sfGFP* carrying pBBR P*_pdc_*-*popZ* under regular growth conditions (OD600 of 0.5 - 0.9). A focus of light phase contrast was observed at one of the cell poles within cells. Scale bar, 5 μm.

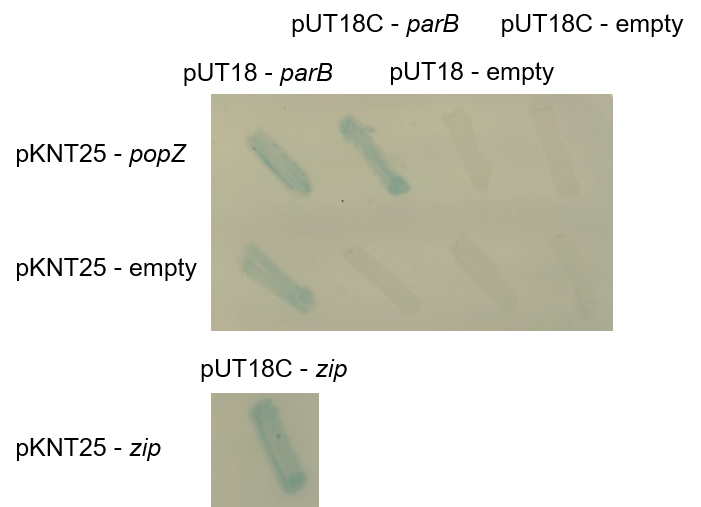

**Fig. S5 *Z. mobilis* ParB appears to interact with *Z. mobilis* PopZ.**

Bacterial two‐hybrid assay by *E. coli* BTH101 expressing pKNT25-popZ and pUT18(C)‐*parB* to examine interaction between PopZ and ParB. As negative controls, pKNT25-empty and pUT18/18C-empty were included in the assay, and for a positive control pKNT25-zip and pUT18C-Zip were co-expressed. The *E. coli* strains were grown on LB agar plate supplemented with X-gal (40 μg/ml) and IPTG (0.5 mM) for 24 hours, and the images were taken. The blue colour indicates the interaction between fusion proteins because it reconstitutes adenylate cyclase activity and activates *lacZ* expression, which yields blue colour on the plates.

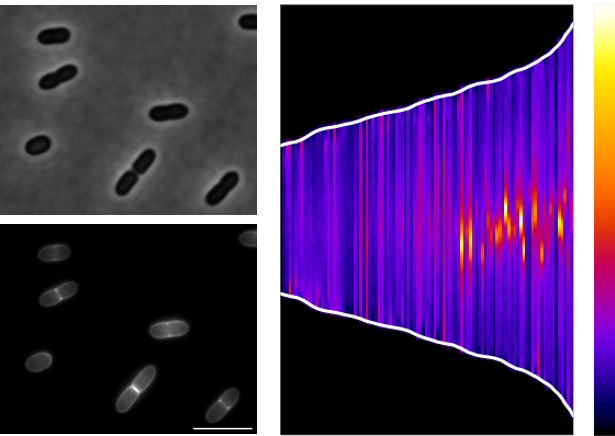

**Fig. S6 Newly inserted PG are localised around the mid-cell in *Z. mobilis***.

(Left) Representative phase contrast (top) and fluorescence (middle) image of growing *Z. mobilis* cells stained by HADA for 20 minutes and spotted on the RM agarose-pad before imaging. Scale bar, 5 μm. (Right): A demograph showing the localization and intensities of HADA fluorescence signal from individual cells in the population. The fluorescence intensity ranged from 1,000 to 10,000, corresponding to the color scale from dark blue to yellow. N = 96. Biological replicates = 2.

**Supplemental tables**

**Table S1.** Bacterial strains and plasmids used in this study.

| Bacterial strains |  |  | Description / Reference |
| --- | --- | --- | --- |
| *Zymomonas mobilis* ZM6 | |  | ATCC29191 purchased from DSMZ |
| *Escherichia coli* DH5α |  |  | Cloning strain (Lab stock) |
| *Escherichia coli* WM6026 | |  | Conjugation strain (1) |
| *Escherichia coli* BTH101 | |  | Strain for bacterial two-hybrid assay (3) |
| *Zymomonas mobilis* + SacB | |  | *B. subtilis* sacB at ZZ*6*_*1449* (2) |
| *Zymomonas mobilis* Δ*popZ* | |  | Δ*ZZ6_1271* (2) |
| *Zymomonas mobilis* Δ*popZ + popZ* | | | Δ*ZZ6_1271* + *ZZ6_1271* (this study) |
| *Zymomonas mobilis parB-sfGFP* | |  | *ZZ6_1203-GFPsf* |
| *Zymomonas mobilis* Δ*popZ* + *parB-sfGFP* | | | *ΔZZ6_1271 + ZZ6_1203-sfGFP* (this study) |
| *Zymomonas mobilis* Δ*popZ + popZ + parB-sfGFP* | | | Δ*ZZ6_1271* + *ZZ6_1271, ZZ6_1203-GFPsf* (this study) |
| *Zymomonas mobilis* pBBR P*pdc-popZ* | | | pBBR P*pdc*-*popZ* (this study) |
| *Zymomonas mobilis pBBR* P*pdc-popZ + parB-sfGFP* | | | *ZZ6_1203-sfGFP*, pBBR P_pdc_-*popZ* (this study) |
| *Escherichia coli BTH101* pUT18, pKNT25 | | | pUT18, pKNT25 (lab stock) |
| *Escherichia coli BTH101* pUT18C, pKNT25 | | | pUT18C, pKNT25 (lab stock) |
| *Escherichia coli BTH101* pUT18 *parB,* pKNT25 | | | pUT18 *parB*, pKNT25 (this study) |
| *Escherichia coli BTH101* pUT18C *parB,* pKNT25 | | | pUT18C *parB*, pKNT25 (this study) |
| *Escherichia coli BTH101* pUT18*,* pKNT25 *popZ* | | | pUT18, pKNT25 *popZ* (this study) |
| *Escherichia coli BTH101* pUT18C*,* pKNT25 *popZ* | | | pUT18C, pKNT25 *popZ* (this study) |
| *Escherichia coli BTH101* pUT18 *parB,* pKNT25 *popZ* | | | pUT18 *parB*, pKNT25 *popZ* (this study) |
| *Escherichia coli BTH101* pUT18C *parB,* pKNT25 *popZ* | | | pUT18C *parB*, pKNT25 *popZ* (this study) |
| *Escherichia coli BTH101* pUT18C *zip*, pKNT25 *zip* | | | pUT18C *zip*, pKNT25 *zip* (lab stock) |

**Table S2.** Oligonucleotides used in this study.

| Plasmids |  |  |  | Description / Reference |
| --- | --- | --- | --- | --- |
| pPK15534 |  |  |  | Suicide vector (1) |
| pPK15534 *+ sacB* | |  |  | pPK15534 carrying *B. subtilis sacB* (this study) |
| pPK15534 *+ sacB + parB-sfGFP* | | |  | pPK15534 + *sacB* carrying *ZZ6_1203-sfGFP* (2) |
| pPK15534 *+ sacB ΔpopZ* | | |  | pPK15534 + *sacB* carrying Δ*ZZ6_1271* cassettes (2) |
| pUT18 |  |  |  | Plasmid used for Bacterial two-hybrid assay (3) |
| pUT18C |  |  |  | Plasmid used for Bacterial two-hybrid assay (3) |
| pUT18 *parB* |  |  |  | pUT18 carrying *ZZ6_1203* (this study) |
| pUT18C *parB* | |  |  | pUT18C carrying *ZZ6_1203* (this study) |
| pKNT25 *popZ* | |  |  | pKNT25 carrying *ZZ6_1271* (this study) |
| pUT18C zip |  |  |  | pUT18C carrying *zip* used for positive control (3) |
| pKNT25 *zip* |  |  |  | pKNT25 carrying zip used for positive control (3) |
| pPK15534 *+ sacB + popZ* | | |  | pPK15534 + *sacB* carrying *ZZ6_1271* (this study) |
| pBBR *+ pdc-popZ* | |  |  | pBBR carrying P*pdc*-*ZZ6_1271* (this study) |
| References: | (1) Lal et al, Front. Microbiol. 10:2216. doi: 10.3389/fmicb.2019.02216 | | | |
|  | (2) Fuchino et al, bioRxiv doi:10.64898/2026.01.13.699301:2026.01.13.699301. | | | |
|  | (3) Karimova G et al, Proceedings of the National Academy of Sciences 95:5752-5756. | | | |

**Table S2.** Oligonucleotides used in this study.

| Oligo | Sequence | Use |
| --- | --- | --- |
| **NKF99** | acccgtggttcatgcatcagcgtatggggctgacttcaggtgc | ***popZ complementation*** |
| **NKF100** | acgccttttctagcaaaggggtaattctcatgtttgacagcttatcac |  |
| **NKF219** | cacctgaagtcagccccatacgctgatgcatgaaccacgggtg |  |
| **NKF224** | gctgtcaaacatgagaattacccctttgctagaaaaggcgtgcc |  |
| **NKF378** | gcttcctacattcttggaaagtcgttttagttatatcttgggcttgctc |  |
| **NKF379** | ccaagatataactaaaacgactttccaagaatgtaggaagcccgac |  |
| **NKF380** | agggggtataatccggtctcattagaaatgctggccagtaattctggag |  |
| **NKF381** | ttactggccagcatttctaatgagaccggattataccccctaggaac |  |
| **NKF340** | aatcatgtctagattcaaggtgtcccgttcctttttccc | ***pBBR Ppdc*-*popZ*** |
| **NKF382** | tttgaatatatggagtaagcaatgcgccctgaaccttcgatgg |  |
| **NKF383** | tcgaaggttcagggcgcattgcttactccatatattcaaaacactatgtc |  |
| **NKF384** | aatcatgctcgagttagaaatgctggccagtaattctggag |  |
| **NKF391** | aatcatgtctagagcgccctgaaccttcgatggaagac | **pKNT25 *popZ*** |
| **NKF392** | aatcatgggtaccttagaaatgctggccagtaattctggag |  |
| **NKF393** | aatcatgtctagagagtttcgacaagaaaaaccgtcctcg | **pUT18 *parB*** |
| **NKF394** | aatcatgggtaccttagaaatcagagccggaaagccg |  |
| **NKF395** | aatcatgaagcttgagtttcgacaagaaaaaccgtcctcg | **pUT18C *parB*** |
| **NKF396** | aatcatgtctagaccgaaatcagagccggaaagccgttg |  |
